# Choosing between cocaine and sucrose under the influence: testing the effect of cocaine tolerance

**DOI:** 10.1101/2021.04.02.438165

**Authors:** Y. Vandaele, S.H Ahmed

## Abstract

Cocaine use not only depends on the reinforcing properties of the drug, but also on its intoxicating effects on alternative nondrug activities. In animal models investigating choice between cocaine and alternative sweet rewards, the latter influence can have a dramatic impact on choice outcomes. When cocaine intoxication at the moment of choice is prevented by imposing sufficiently long intervals between choice trials, animals typically prefer the sweet reward. However, when choosing under the drug influence is permitted, animals shift their preference in favor of cocaine. We previously hypothesized that this preference shift is mainly due to a direct suppression of responding for sweet reward by cocaine intoxication. Here we tested this hypothesis by making rats tolerant to this drug-induced behavioral suppression. Contrary to our expectation, tolerance did not prevent rats from shifting their preference to cocaine when choosing under the influence. Thus, other mechanisms must be invoked to explain the influence of cocaine intoxication on choice outcomes.

## Introduction

Whether recreational or problematic, drug use not only depends on the inherent reinforcing properties of the drug but also depends on its pharmacological or intoxicating effects on alternative behavioral activities. Some of these effects can be beneficial for other nondrug-related behaviors (Müller, 2020; Müller and Schumann, 2011; Pickard, 2020). For instance, psychostimulants are often consumed to improve social interactions, to enhance cognitive performance or to counteract fatigue (Müller and Schumann, 2011). Alternatively, other drug effects can be detrimental to alternative nondrug activities, notably in the context of substance use disorder (SUD). Drug interference with alternative activities may contribute, at least partly, to explain some diagnostic criteria of SUD in the Diagnostic and Statistical Manual of Mental Disorders (DSM, 5^th^ edition), including: (1) “Substance use has caused relationship problems or conflicts with others”, (2) “Failure to meet responsibilities at work, school, or home because of substance use” and (3) “Activities are given up in order to use the substance” (American Psychiatric Association, 2013). The suppression of non-drug related alternatives by drug use can create a vicious circle where the drug becomes the most valuable option available (Heyman, 2010). On the other hand, drug interference with important non-drug related activities can motivate the decision to abstain and could constitute a strong incentive for addiction recovery (Branch, 2011; Heyman, 2013, 2010). Thus, investigating how drug effects can influence non-drug related behaviors is important when considering the transition to or recovery from SUD.

In animal models of SUD involving drug self-administration in presence of alternative nondrug rewards, choosing under the influence of the drug is typically prevented by imposing sufficiently long inter-trial intervals (ITI), allowing for drug dissipation between choice trials (Cantin et al., 2010; Lenoir et al., 2013a, 2007). In this condition, the large majority of rats prefer the alternative nondrug reward. This finding, first discovered more than 10 years ago (Lenoir et al., 2007), was repeatedly reproduced in a large set of conditions, including: different drug and nondrug rewards; various drug doses and history of drug self-administration; different reward delays and costs (Augier et al., 2012; Cantin et al., 2010; Caprioli et al., 2015; Huynh et al., 2017; Kearns et al., 2017; Kerstetter et al., 2012; Lenoir et al., 2013b; Lenoir and Ahmed, 2008; Madsen and Ahmed, 2015; Pelloux and Baunez, 2017; Russo et al., 2018; Schwartz et al., 2017; Venniro et al., 2018). However, in a free-operant choice schedule in which both options are continuously available, and thus, in which choosing under the drug influence is permitted, choice behavior dramatically differs (Bozarth and Wise, 1985; Freese et al., 2018; Thomsen et al., 2013, 2008; Vandaele et al., 2016). Notably, rats offered a choice between cocaine and saccharin in these conditions first self-administer saccharin before switching to cocaine exclusively until the end of the session. A similar choice pattern was observed in a discrete-trial schedule when the ITI is sufficiently shortened to permit choice under the influence (Kerstetter et al., 2012; Vandaele et al., 2016). Finally, when the drug influence is induced artificially before each choice trial by a non-contingent injection of cocaine, this is sufficient to bias choice toward cocaine (Freese et al., 2018; Guillem and Ahmed, 2018; Vandaele et al., 2016). Thus, when rats are choosing between cocaine and saccharin under the influence of cocaine, they shift their preference to cocaine. The mechanisms underlying this drug-induced shift in preference are yet to be fully understood.

We have previously suggested that cocaine intoxication shifts choice to cocaine through direct suppression of the alternative nondrug-related behavior. First, as a psychostimulant, cocaine exerts potent anorexic effects, suppressing both feeding and drinking behaviors in rats (Balopole et al., 1979; Cooper and Francis, 1993; Vandaele et al., 2016; Wolgin and Hertz, 1995; Woolverton et al., 1978). Second, cocaine-induced suppression of responding for saccharin is strongly correlated with cocaine-induced shift to cocaine choice (Vandaele et al., 2016). Furthermore, cocaine-induced shift in preference is associated with a suppression of the activity of saccharin-coding neurons in the orbitofrontal cortex and no facilitation of the activity of cocaine-coding neurons (Guillem and Ahmed, 2018). Finally, drug-induced shift in preference is not observed with heroin intoxication which is known to enhance rather than suppress eating and drinking behaviors (Cooper et al., 1985; Parker et al., 1992; Vandaele et al., 2016; but see Chow and Beckmann, 2021; Townsend et al., 2021). Thus, the comparison of cocaine and heroin in a free-operant choice schedule or following pre-trial drug injections suggests that their opposite effect on preference is likely mediated by their opposite effects (suppressing versus enhancing) on the alternative nondrug reward, rather than by their common priming effects on drug seeking, as seen in other behavioral paradigms (Ahmed and Cador, 2006; de Wit and Stewart, 1983).

The goal of this study is to test more directly the role of the suppressive effects of cocaine in biasing choice in favor of exclusive drug use in a free-operant setting. To this end, tolerance to these effects was induced before choice testing. Previous research showed that tolerance can be induced in hungry rats when amphetamine or cocaine is administered immediately before access to food (Wolgin, 2000; Wolgin and Hughes, 1997; Woolverton et al., 1978). This tolerance is called contingent tolerance because its development requires that animals experience the suppressive effects of the drug while they are eating, or at least, trying to (Wolgin, 2000; Wolgin and Jakubow, 2004). Indeed, if animals are no longer under the influence of amphetamine or cocaine when given access to food, all else being equal, they do not develop tolerance. In our conditions, contingent tolerance was induced by allowing hungry rats to self-administer cocaine immediately before a short access to sucrose-sweetened water. If cocaine intoxication shifts choice in favor of drug use by suppressing responding for sucrose, then tolerance to these suppressive effects should prevent or retard such preference shift.

## Materials and methods

### Subjects

Twenty-eight male Wistar rats weighting in average 275-300g at the beginning of the experiment were used (Charles River, L’Arbresle, France). Rats were housed in groups of 2 in a temperature-and light-controlled vivarium (21°C, reversed 12-h light-dark cycle). Rats were food-restricted and maintained at 80% of their estimated free-feeding weight. Water was freely available in the home cages during behavioral testing. Three rats were excluded due to failure in catheter patency, leaving a total of 25 rats for the analysis. All experiments were conducted in accordance with institutional and international standards of care and use of laboratory animals (UK Animals (Scientific Procedures) Act, 1986; and associated guidelines; the European Communities Council Directive (2010/63/UE, 22 September 2010) and the French Directives concerning the use of laboratory animals (décret 2013-118, 1 February 2013). The animal studies were reviewed and approved by the Committee of the Veterinary Services Gironde, agreement number B33-063-5.

### Apparatus

Fourteen identical operant chambers (30 x 40 x 36 cm) described in detail elsewhere (Lenoir et al., 2013a) were used (Imetronic, Pessac, France). Chambers were equipped with two retractable levers, a commercially-available lickometer circuit, two syringe pumps, a single-channel liquid swivel (Lomir biomedical Inc., Quebec, Canada) and two pairs of infrared beams to measure locomotor activity.

### Surgery

Rats received a surgery for the implantation of chronic silastic catheters (Dow Corning Corporation, Michigan, USA) in the right jugular vein, exiting the skin in the middle of the back about 2 cm below the scapulae, as described previously (Lenoir et al., 2013a).

### Operant training

Animals were first trained to press on the left lever for a solution of 20% sucrose under a fixed-ratio 1 (FR1 time-out 20s) schedule as described in detail elsewhere (Lenoir et al., 2013a). We chose to train hungry animals with a caloric solution of 20% sucrose to increase the stakes for the consumption of the nondrug reward and favor the development of a contingent tolerance to the suppressive effects of cocaine. Discrete volumes of sucrose were delivered in the adjacent drinking cup by voluntary liking over the time-out period of 20s, signaled by the illumination of the cue-light above the lever. The drinking cup was automatically filled with 2 volumes over the first 3-s and additional volumes were obtained by licking, resulting in a maximum volume delivered of 0.32mL. Responses during the 20-s time out were not rewarded. Sessions ended after rats had earned a maximum of 30 rewards or 3h had elapsed. Rats were trained under a FR1 schedule for 6 sessions followed by 3 sessions under a fixed ratio 2 (FR2) schedule. Rats were then trained to self-administer intravenous cocaine (0.25 mg delivered over 5 s) by pressing on the right lever under a FR2 schedule for 10 sessions (Figure 1A). Sessions were limited to a maximum of 3h or 50 injections. Importantly, rats were systematically tethered to the infusion line to equate training conditions across sucrose and cocaine rewards.

**Figure 1:**
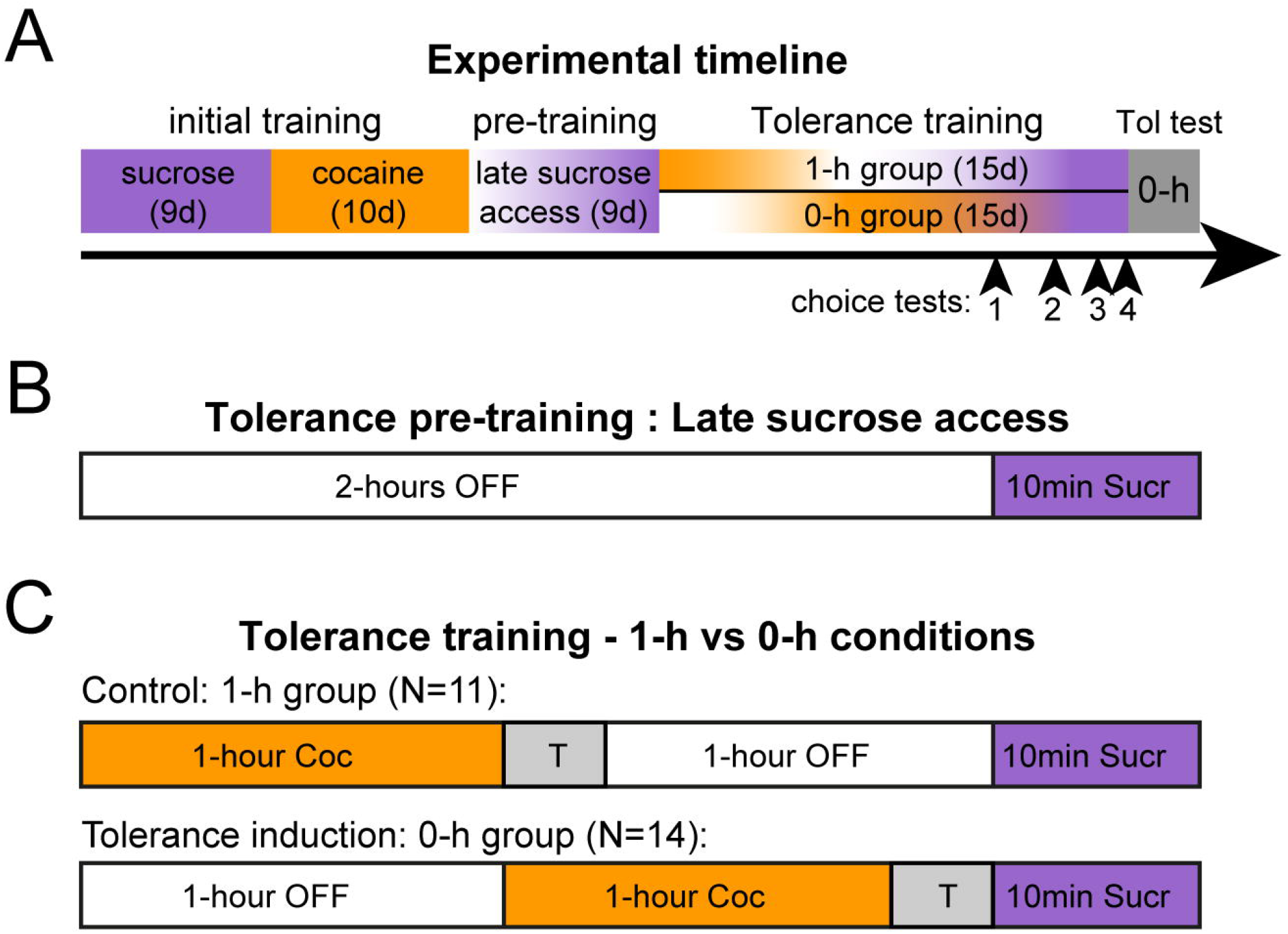
Schematic representation of the procedure. **A.** Experimental timeline. Arrow heads indicate choice tests. **B.** After initial sucrose and cocaine self-administration training, all rats are offered a 10-min sucrose access after a 2-h period with no reward available (OFF). **C.** During tolerance training, one group of rats receive 1-h cocaine access followed by a 1-h break before the 10-min sucrose access (control group, 1-h condition). In the other group, cocaine self-administration occurs after 1-h and immediately before the 10-min sucrose access (tolerant group, 0-h condition). The gray box marked with a ‘T’ represents a transition period during which rats can receive an extra cocaine injection before the next phase.

### Tolerance training procedure

After acquisition of lever pressing for sucrose and cocaine, rats were allowed to self-administer sucrose for 10-minutes under a FR2 schedule after an initial 2-h period with no reward available (Figure 1B). The levers remained retracted during the first 2-h and extension of the left lever signaled the onset of sucrose availability. During the first 3 sessions, half of the rats failed to notice the lever insertion. Thus, the onset of sucrose access was also signaled by the illumination of the house light and one free delivery of 0.08mL of sucrose for three additional sessions. Upon correct acquisition of sucrose self-administration under these conditions, the house light cue and free sucrose delivery were removed and training pursued for three more sessions.

One hour of cocaine self-administration was then introduced either immediately (0-h group; N=14) or 1-h before sucrose access (1-h group; control condition; N=11) (Figure 1C). Thus, only rats in the 0-h group were under the influence of cocaine during sucrose self-administration. To minimize the delay between the last cocaine injection and subsequent access to sucrose in the 0-h group, we introduced a transition period (T) after the cocaine phase that ended immediately after the first cocaine injection or after a maximal duration of 10-min (Figure 1C). Importantly, only in two rare occasions did a rat missed the opportunity for this last cocaine injection. To equate training conditions, this transition period was also introduced at the end of cocaine self-administration in the 1-h group. Rats were randomly assigned to either group.

### Free-operant choice procedure

After acquisition of contingent tolerance in the 0-h group, rats’ preference was tested in the free-operant choice procedure. Both levers were simultaneously presented throughout the duration of the 2-h session. Completion of the FR2 response requirement on either lever resulted in the delivery of the corresponding reward (i.e., intravenous cocaine injection or 20-s sucrose access), signaled by the illumination of the cue light above the selected lever during the 20-s time out. Two variants of this procedure were tested. In the first variant, rats were first allowed to self-administer cocaine for 20-min before initiation of the 2-h choice session (test 2). In the second variant, the duration of the session was extended to 5-h (test 4). Few baseline sessions with the tolerance training procedure were conducted between choice sessions. At the end of choice testing, tolerance expression was tested in all rats during a single tolerance test session conducted in the 0-h condition (Figure 1A).

### Data Analysis

Licking efficiency was calculated by computing the ratio of the number of volumes delivered divided by the number of volumes available, during every sucrose accesses. All data were subjected to mixed analyses of variance (ANOVA), followed by post-hoc comparisons using the Tukey’s Honestly Significant Difference (HSD) test, when appropriate. Comparisons with a fixed theoretical level (e.g., 50%) were conducted using one sample t-tests. Some behavioral variables did not follow a normal distribution and were thus analyzed using non-parametric statistics (i.e., Friedman’s test for main effect followed by Wilcoxon’s test for paired comparisons; Mann Whitney for group comparison).

## Results

### Development of a tolerance to cocaine suppressive effects in the 0-h group

During tolerance training, both groups of rats self-administered cocaine similarly during cocaine access at the beginning of the session, 1-h before sucrose access (1-h group) or immediately before (0-h group) (Figure 2A; Mann Whitney test, Z-values <0.74, p-values>0.4). As expected, responding for sucrose was drastically suppressed in rats from the 0-h group, intoxicated with cocaine during sucrose access (group 0-h; Figure 2B). Rats in this group earned significantly less sucrose rewards during the first tolerance training session compared to baseline with a 77.1±4.5% suppression (Figure 2A; Wilcoxon test, Z14=3.30; p<0.001). However, with repeated sessions in the tolerance training procedure, most 0-h rats learned to resist to the suppressive effects of cocaine and progressively increased their responding for sucrose despite cocaine intoxication (Figure 2B, group 0-h (tol): N=10; Friedman test, ANOVA Chi-sqr=48.21; p<0.0001). This subgroup of 10 rats did not differ from 1-h rats during the last three tolerance training sessions (Figure 2B; Mann Whitney test, Z-values <1.73, p-values>0.05). However, sucrose self-administration remained significantly suppressed across tolerance training sessions in 4 rats from the 0-h group, compared to the 1-h group (Figure 2B; 0-h (no tol): N=4; Mann Whitney test, Z-values >2.68, p-values<0.01 except on the 6^th^ session). These rats kept suppressing sucrose self-administration at 50% of their baseline and thus, did not develop reliable tolerance to the suppressive effects of cocaine.

**Figure 2:**
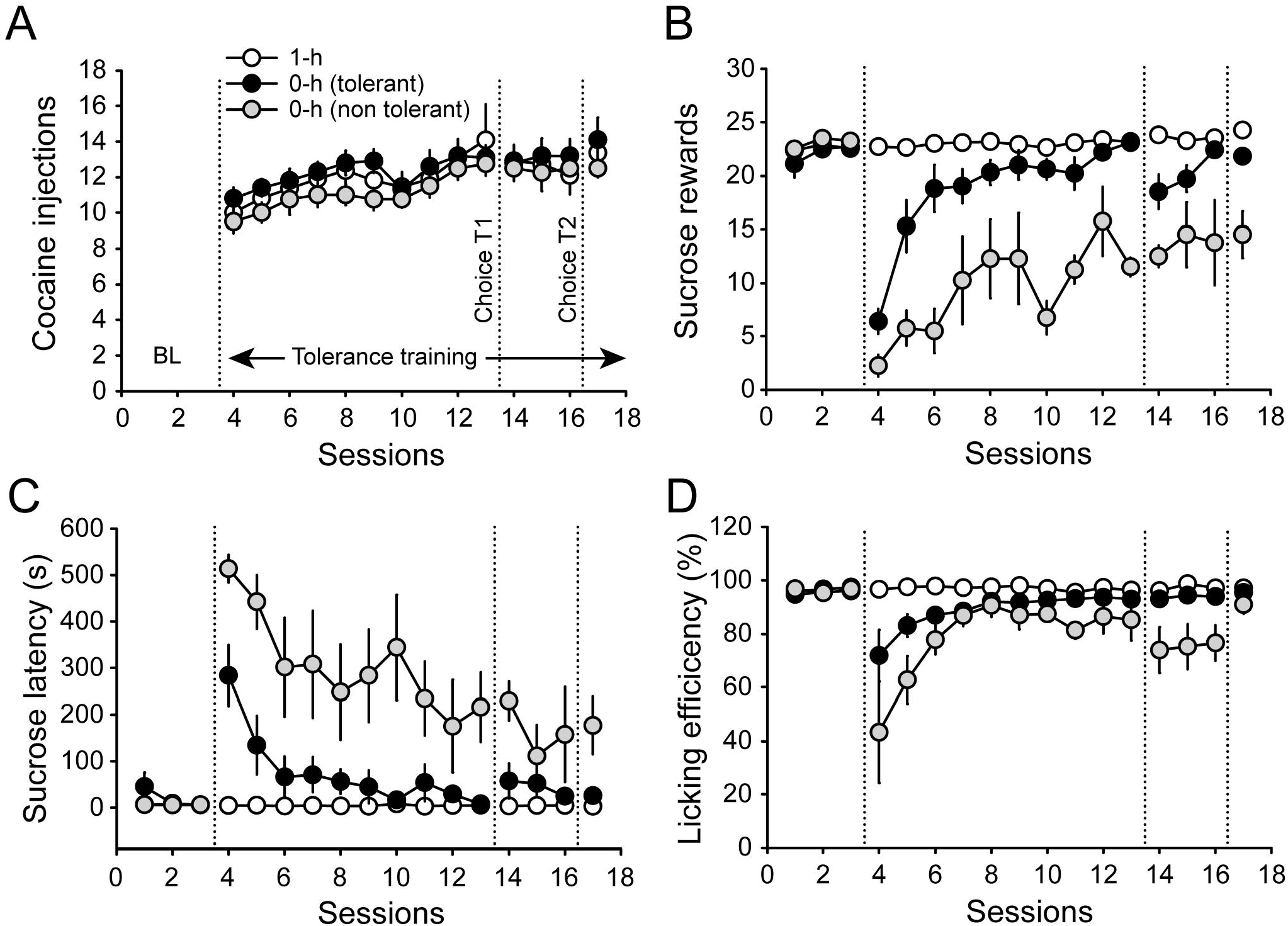
The majority of rats in the 0-h group developed a tolerance to cocaine suppressive effects. A-D. Mean (±SEM) number of cocaine injections (**A**), number of sucrose rewards (**B**), latency to initiate sucrose self-administration (**C**) and licking efficiency (**D**) across tolerance training sessions, in the 1-h group (white circles) and in tolerant rats (black circles) or non-tolerant rats (gray circles) of the 0-h group. Vertical dotted lines delimit tolerance training onset and mark the time of the first and second choice tests (Choice T1 and T2).

Analysis of the latency to initiate sucrose self-administration reveals that the first sucrose access was significantly delayed in 0-h rats compared to 1-h rats during the first session (Figure 2C; Mann Whitney test, Z=3.61, p<0.0001). Rats developing a tolerance to cocaine suppressive effects progressively learned to respond for sucrose from the onset of sucrose availability (Friedman test, ANOVA Chi-sqr=45.02, p<0.0001), with a latency comparable to the 1-h group during the last session (Mann Whitney, Z=−1.76, p>0.05). However, rats failing to develop tolerance maintained a significant post-cocaine delay to initiate sucrose self-administration compared to the 1-h group (Figure 2C; last session: Mann Whitney, Z=−2.22, p<0.05).

Cocaine intoxication during sucrose access not only delayed initiation of sucrose self-administration, and consequently, the number of sucrose rewards, but also interfered with licking behavior as evidenced by a decrease in licking efficiency in the 0-h group compared to baseline (Wilcoxon test, Z=3.23, p<0.01) and compared to the 1-h group (Figure 2D; First session: Mann Whitney, Z=3.45, p<0.001). This result indicates that rats in the 0-h group did not consume all the volumes available during sucrose accesses. However, rats developing a tolerance learned to overcome this suppressive effect on licking behavior and reached a similar licking efficiency as the 1-h group during the last session (Figure 2D; Mann Whitney: Z=1.16, p>0.2). Licking efficiency also increased in 0-h non-tolerant rats but never reached the level of 1-h rats (Figure 2D). Overall, the four 0-h non tolerant rats maintained a clear suppression of sucrose self-administration despite tolerance training. Since our approach is to assess the effect of tolerance to the suppressive effects of cocaine on preference, these rats were excluded from the group 0-h. Furthermore, the low number of 0-h non tolerant rats (N=4) precludes their inclusion in statistical analyses. Thus, we analyzed individual choice patterns in subsequent tests for these rats, separately (suppl figure 1).

### Tolerance to cocaine suppressive effects did not prevent cocaine-biased shift in preference

Preference between cocaine and sucrose was first tested during a 2-h free-operant choice session. Surprisingly, the percentage of cocaine choice did not significantly differ between groups despite a trend toward lower preference for cocaine in 0-h rats (Figure 3A; t_19_=2.05; p=0.054). In fact, the main pattern of choice was similar between groups; all rats first began self-administering sucrose before shifting to cocaine, with no group difference in this first phase (Figure 3B; t_19_=−0.83; p>0.4). Then, most rats continued to self-administer cocaine exclusively until the end of the session, but some occasionally sampled the sucrose option before switching back to cocaine (Figure 2E-F). To quantify this behavior, we assessed the number of inter-reward transitions and found that tolerant rats in the group 0-h made significantly more transitions than 1-h rats (Figure 3C; Mann Whitney: Z=−2.04; p<0.05). However, analysis of the within-session time course of sucrose accesses revealed no group difference (F_1,19_=0.85, p>0.3) nor group by time bin interaction (Figure 3D; F_11,209_=0.38, p>0.5). Rats that did not develop a tolerance in the 0-h group displayed a choice pattern similar to 1-h rats (supplemental Figure 1A).

**Figure 3:**
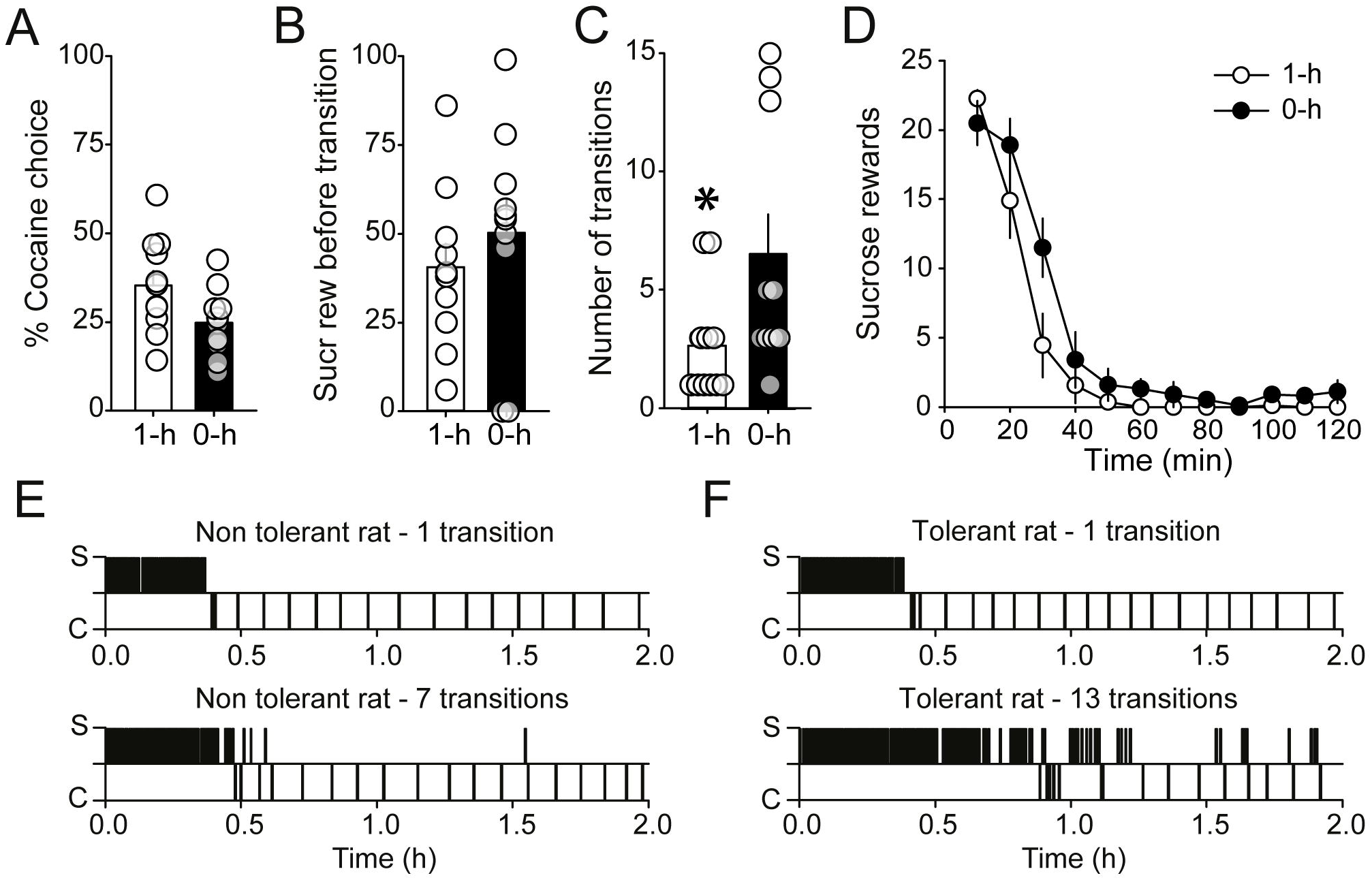
Tolerance to the suppressive effects of cocaine did not prevent cocaine-biased shift in preference. A-C. Mean (±SEM) percentage of cocaine choice (**A**), number of sucrose rewards before transition to cocaine (**B**) and number of inter-reward transitions (**C**) in 1-h and 0-h rats. *p<0.05. **D**. Within-session time course of sucrose rewards in the 1-h and 0-h groups across 10-min time bins. **E-F**. Choice patterns of representative non tolerant rats in the 1-h group (**E**) and tolerant rats in the 0-h group (**F**). Vertical bars above or below the horizontal line represent sucrose (S) and cocaine (C) choices, respectively. For each rat, the number of interreward transitions is indicated.

### Tolerance to cocaine suppressive effects can favor sucrose preference in conditions of high motivation

The results above suggest that although tolerance to the suppressive effects of cocaine allowed some rats to sample sucrose once intoxicated, their usual pattern of sucrose self-administration was prevented. In fact, it seems that although possible, expressing a tolerance to cocaine suppressing effect is difficult. Thus, the initial loading period of sucrose self-administration could be sufficient to reduce sucrose value by sensory-specific satiety, thereby dampening motivation to overcome cocaine suppressive effects, once intoxicated. After three baseline tolerance training sessions, rats were tested in a modified choice session, comprising a 20-min period of exclusive cocaine self-administration, immediately before the 2-h choice session (Choice test 2). We observed no group difference in the number of cocaine injections during the initial 20-min period (Figure 4A; t_19_=0.24, p>0.5). However, preventing the initial sucrose self-administration loading by intoxicating rats from the beginning of the session revealed a significant difference in preference between 0-h and 1-h rats (Figure 4B; Mann-Whitney: Z=2.46, p<0.05). The high preference for cocaine in 1-h rats can be explained by the low number of inter-reward transitions in this group compared to 0-h rats (Figure 4C & E, Mann Whitney: Z=−3.17, p<0.01). In contrast, tolerant rats in the 0-h group succeeded to make at least one inter-reward transition to self-administer sucrose, generally at the beginning of the choice session (Figure 4C-D & F). Thus, analysis of the within-session time course of sucrose accesses revealed a group difference (F_1,19_=12.20, p<0.01), specifically during the first 10 minutes of the session (Figure 4D; group by time interaction: F_11,209_=2.33, p<0.05; post-hoc 0-10min: p<0.05). Rats in the 0-h group that failed to develop a tolerance expressed choice patterns more comparable to the 1-h group (supplemental Figure 1B).

**Figure 4:**
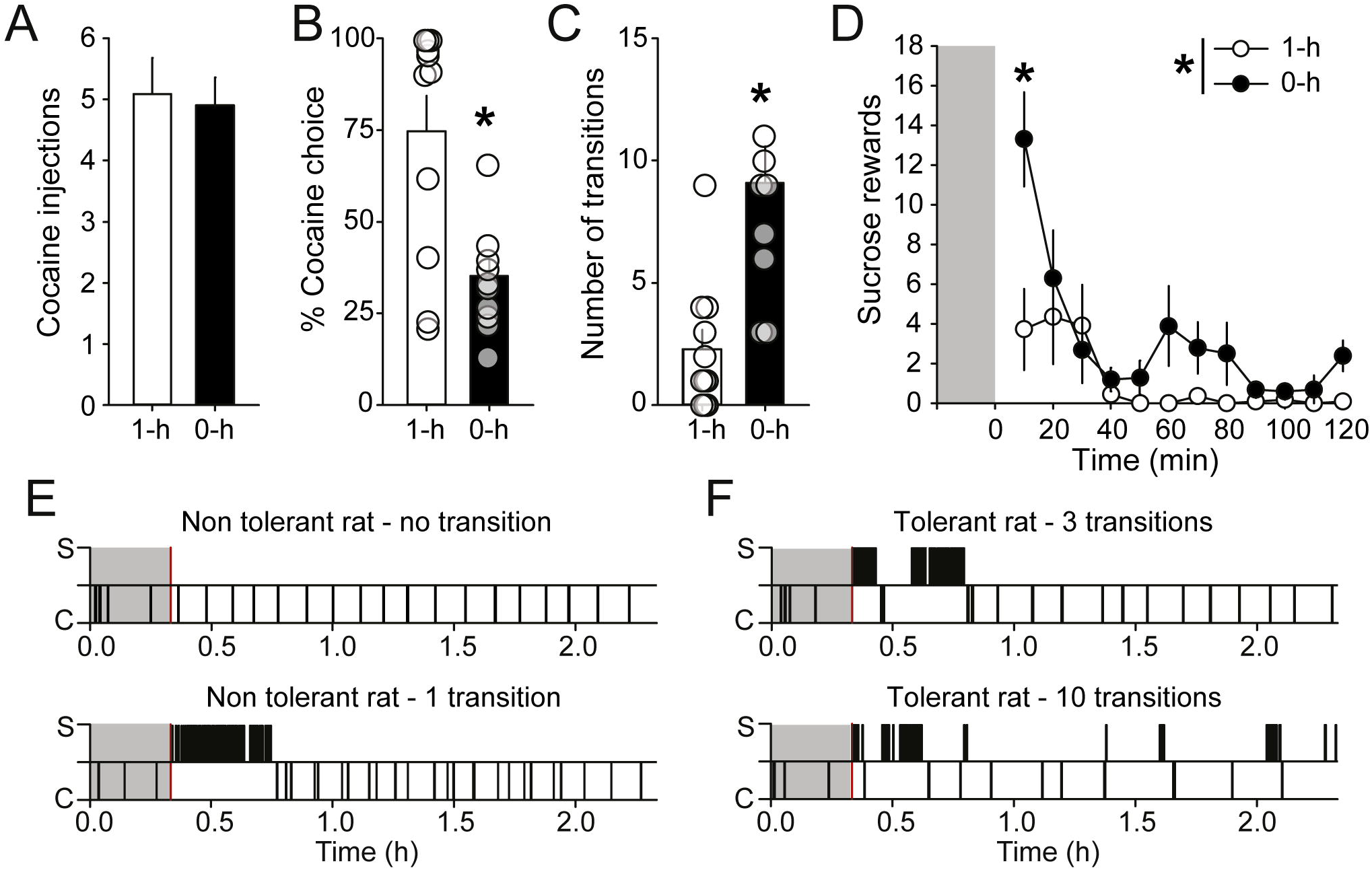
An effect of tolerance to cocaine suppressive effects is revealed when motivation for sucrose is high. **A**. Mean (±SEM) number of cocaine injections during pre-choice cocaine self-administration in 1-h and 0-h rats. (**B-C**) Mean (±SEM) percentage of cocaine choice (**B**) and number of inter-reward transitions (**C**) in 1-h and 0-h rats. * p<0.05. **D**. Within-session time course of sucrose rewards in the 1-h and 0-h rats across 10-min time bins. *p<0.05. **E-F.** Choice patterns of representative non tolerant rats in the 1-h group (**E**) and tolerant rats in the 0-h group (**F**). Vertical bars above or below the horizontal line represent sucrose (S) and cocaine (C) choices, respectively. For each rat, the number of inter-reward transitions is indicated. In figures D-F, the gray area represents the 20-min period of pre-choice cocaine self-administration. The onset of the choice session is marked with a red vertical bar in E-F.

These results suggest that when their motivation for sucrose was sufficient, 0-h rats expressed a tolerance to the suppressive effects of cocaine and maintained a preference for sucrose despite prior cocaine intoxication (t-test against indifference: t_10_=−3.55, p<0.01). Yet, the expression of tolerance was not perfect and sucrose self-administration remained relatively suppressed in comparison to tolerance training sessions in which sucrose is the only reward available. Importantly, group differences in choice behavior disappeared when rats were tested for a second time in a 2-h choice session, after one baseline session, in absence of prior cocaine self-administration as in the first choice session (Supplementary figure 2; % cocaine choice; Mann Whitney: Z=1.26, p>0.2).

### Tolerance to cocaine suppressive effects did not favor sucrose choice during an extended 5-h choice session

We next asked whether given sufficient time, rats tolerant to cocaine suppressive effects would eventually switch back to sucrose after the shift to cocaine choices. Rats were tested in the final choice session for a duration of 5-h. Increasing the session duration had no effect on the expression of tolerance during choice behavior. There was no group difference in the percentage of cocaine choice (Figure 5A; t_19_=1.59, p>0.1) or in the number of sucrose access before the first transition (Figure 5B; t_19_=0.71, p>0.7). Although some rats in both groups made a high number of inter-reward transitions, the groups did not differ on this variable (Figure 5C; Mann Whitney: Z=−1.16, p>0.2). Accordingly, we observed no group difference in the within-session time course of sucrose accesses (Figure 5D; F_1,19_=2.58, p>0.1).

**Figure 5:**
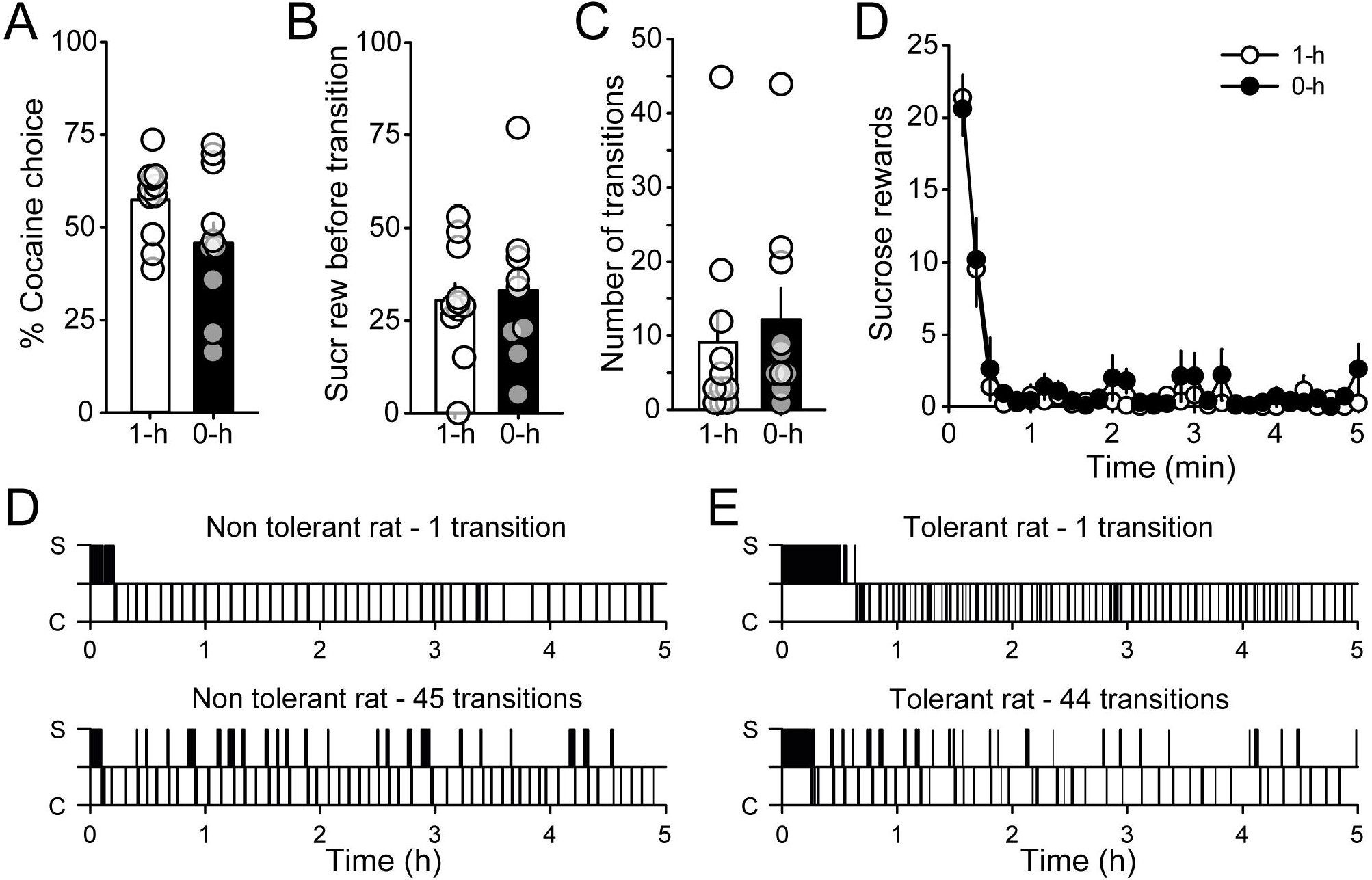
Tolerance to cocaine suppressive effects had no effect on preference during an extended 5-h choice session. **A-C.** Mean (±SEM) percentage of cocaine choice (**A**), number of sucrose rewards before transition to cocaine (**B**) and number of inter-reward transitions (**C**) in 1-h and 0-h rats. **D.** Within-session time course of sucrose rewards in the 1-h and 0-h rats across 10-min time bins. **E-F.** Choice patterns of representative non tolerant rats in the 1-h group (**E**) and tolerant rats in the 0-h group (**F**). Vertical bars above or below the horizontal line represent sucrose (S) and cocaine (C) choices, respectively. For each rat, the number of interreward transitions is indicated.

Few rats in the 1-h group made a high number of inter-reward transitions (i.e. 45 transitions, Figure 5D, bottom panel). Alternatively, some tolerant rats made only a few numbers of inter-reward transitions (Figure 5E, top panel). These results suggest that repeated choice testing may have favored the development or altered the expression of tolerance in 1-h and 0-h rats, respectively. To test this hypothesis, all rats were tested in a tolerance training session and allowed to self-administer cocaine for 1-h immediately before sucrose access (0-h condition). Cocaine and sucrose self-administration during this test was compared between groups and within-subject with respect to the last baseline session, conducted with the respective 0-h and 1-h tolerance training conditions.

Rats reliably self-administered cocaine with no difference between groups and compared to the baseline session (Figure 6A; effect of group: F_1,19_=0.16, p>0.6; effect of session: F_1,19_=0.07, p>0.7). Prior cocaine self-administration significantly suppressed sucrose self-administration in both groups compared to baseline (Figure 6B; effect of session: F_1,19_=75,4, p<0.0001), suggesting that tolerance was partially lost in 0-h rats. However, there was a significant session by group interaction (Figure 6B; F_1,19_=8.62, p<0.01). Post-hoc analysis reveals that suppression of sucrose self-administration was stronger in the 1-h group compared to the 0-h group (Figure 6B; p<0.05). Analysis of the latency to initiate sucrose self-administration reveals a significant group by session interaction (Figure 6C; F_1,19_=20.05, p<0.001). Although the number of sucrose rewards earned by 0-h rats was lower than baseline, these rats initiated sucrose self-administration with a comparably short latency (Figure 6C; post-hoc p>0.9). In sharp contrast, sucrose self-administration was considerably delayed in rats from the 1-h group compared to baseline (Figure 6C; post hoc: p<0.001). Cocaine intoxication altered licking efficiency in both groups of rats, with no significant group by session interaction (Figure 6D; effect of session F_1,19_=13.31, p<0.01). Together these results suggest that rats in the 1-h group did not develop a tolerance to cocaine suppressive effects with repeated choice testing. However, rats in the 0-h group partially lost their tolerance. Interestingly, the expression of tolerance during the test, assessed by the number of sucrose rewards earned under the influence of cocaine, was positively correlated with the number of inter-reward transitions during the preceding 5-h choice session (Figure 6E; r=0.51, p<0.05).

**Figure 6:**
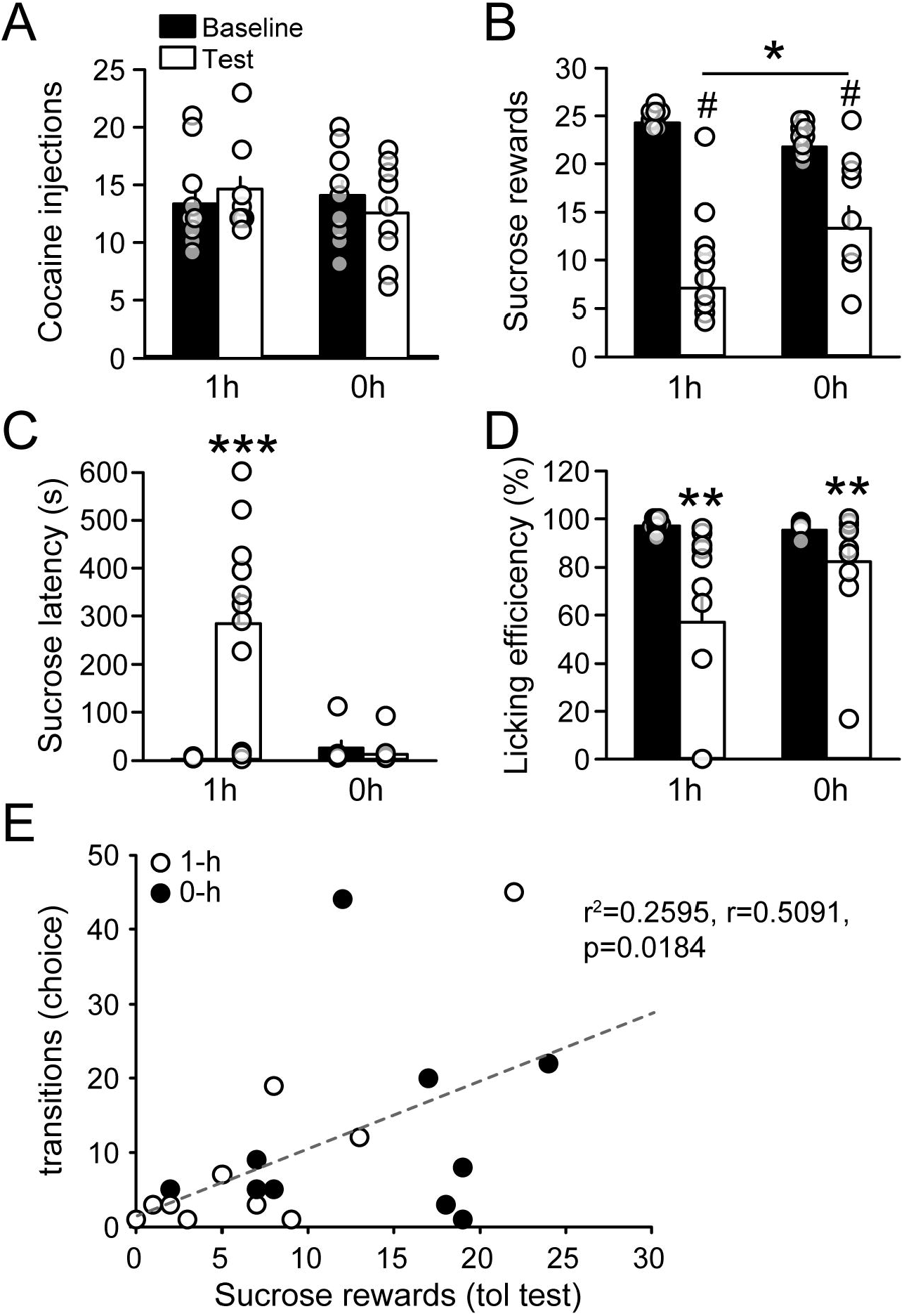
Partial loss of tolerance in 0-h rats with repeated choice sessions. **A-D.** Mean (±SEM) number of cocaine injections (**A**), number of sucrose rewards (**B**), latency to initiate sucrose self-administration (**C**) and licking efficiency (**D**) during the tolerance test session (white bars) compared to baseline (black bars), in 1-h and 0-h rats. Baseline refers to the last tolerance training session. *p<0.05, **p<0.01, ***p<0.001, #p<0.0001. **E.** Correlation between the number of sucrose rewards earned during the tolerance test and the number of inter-reward transitions during the preceding 5-h choice session. White and black circles represent rats from the 1-h and 0-h groups, respectively.

## Discussion

The majority of rats exposed to sucrose while intoxicated learned to tolerate the suppressive effects of cocaine on responding for sucrose, confirming and extending previous research (Wolgin, 2000; Wolgin and Hertz, 1995; Wolgin and Jakubow, 2004). However, this tolerance only had a small effect on preference during subsequent choice under the influence. As reported previously, during free-operant choice, non-tolerant rats first chose sucrose before eventually switching to cocaine nearly exclusively until the end of the session. Overall, the same behavior was also observed in tolerant rats, except that once intoxicated, they tended to transition more between the two rewards. A significant effect of tolerance was only manifest when rats were under the influence of cocaine before onset of the choice session. Thus, contrary to our expectation, tolerance did not prevent rats from shifting their preference to cocaine when choosing under the influence. Other mechanisms must be invoked to explain the influence of cocaine intoxication on choice outcomes in free-operant choice schedule.

By the end of tolerance training, rats in the 0-h group expressed a robust tolerance to cocaine suppressive effects, self-administering sucrose with the same performance and efficiency as the control group. As predicted by Wolgin et al (Wolgin, 2000; Wolgin and Hertz, 1995; Wolgin and Jakubow, 2004), this tolerance only developed when rats had access to sucrose while intoxicated by cocaine. Although we did not observe tolerance learning in 1-h rats across repeated choice sessions, a partial extinction of tolerance occurred in some 0-h rats, in agreement with prior research (Woolverton et al., 1978). Importantly, the expression of tolerance at the end of choice testing was correlated with the number of inter-reward transitions during choice, suggesting that the suppressive effects of cocaine on responding for sucrose somehow influenced choice behavior. However, in most choice sessions, the learned tolerance to cocaine suppressive effects did not generalize well to the free-operant choice procedure.

In the present study, the lack of generalization of tolerance to the choice setting could be explained by several non-exclusive hypotheses. The contingent tolerance to cocaine suppressive effects is commonly described as a form of instrumental learning, and as such, is context-dependent (Wolgin, 2000). Thus, it is possible that, although rats were trained and tested in the same conditioning chambers, the settings for tolerance training and choice testing were still considered as distinct instrumental contexts. Another hypothesis is that, by the end of tolerance training, sucrose self-administration was under habitual control, and thus, not sensitive to the current motivation level for sucrose (Vandaele et al., 2020, 2019). In other words, motivation for sucrose would be suppressed but this suppression would not affect habitual sucrose seeking behavior during tolerance training. Testing preference between cocaine and sucrose in a free-operant choice setting would then reengage goal-directed control (Halbout et al., 2016; Holland, 2004; Kosaki and Dickinson, 2010; Vandaele and Ahmed, 2020), and reveal cocaine suppressing effects on responding for sucrose. Habitual learning during tolerance training could be partly explained by prior cocaine exposure, known to promote habit (Corbit et al., 2014; Gourley et al., 2013; LeBlanc et al., 2013; Miles et al., 2003).

An alternative hypothesis directly supported by the data is that rats would be more motivated by sucrose during tolerance training compared to choice testing. Indeed, during tolerance training, the hungry rats only received 10-min access to sucrose at the end of the session, after a waiting time of 2-h. In contrast, during choice sessions, rats self-administered sucrose continuously from the very beginning of the session, for about 20-30 minutes. During this loading period, one should expect that motivation for sucrose progressively decreases, at least partly, by sensory-specific satiety, a process that should increase the probability of initiating cocaine use and, thus, of the subsequent preference shift. Thus, although rats had learned to tolerate the suppressive effects of cocaine, controlling drug-induced stereotypies is more difficult when the motivation for sucrose has decreased. Supporting this hypothesis, we showed that an effect of tolerance was only manifest when the initial loading period of sucrose self-administration was prevented by intoxicating rats before onset of the choice session. In these conditions, tolerant rats succeeded to resist cocaine suppressive effects and were able to respond for sucrose few times under the influence of cocaine. However, even in these conditions, the effect of tolerance was modest since the pattern of sucrose self-administration was significantly suppressed by cocaine self-administration during choice testing.

Preference during free-operant choice not only depends on motivation for sucrose and expression of tolerance to the suppressive effects of cocaine, but can also depend on the motivation for the drug itself. Indeed, cocaine intoxication can prime responding for cocaine in behavioral paradigms such as drug-induced reinstatement (Ahmed and Cador, 2006; de Wit and Stewart, 1981; Shaham et al., 2003). Thus, in a free-operant setting, cocaine choice can transiently enhance motivation for cocaine by increasing cocaine incentive value (Robinson and Berridge, 1993) or by inducing a negative affective state (i.e. withdrawal) that would be alleviated by another dose of cocaine (Ettenberg, 2004; Koob and Le Moal, 2001). Similar processes can be observed for opioids (de Wit and Stewart, 1983; Shaham et al., 2003). We previously suggested that, in contrast to cocaine, heroin exerts orexigenic effects and that these effects would enhance, rather than suppress, responding for the alternative nondrug reward, when choosing under the influence is permitted (Vandaele et al., 2016). However, recent findings suggest that the processes controlling drug-vs-food choice under opioid influence cannot be limited to the drug orexigenic effects (Chow and Beckmann, 2021; Townsend et al., 2021). Indeed, increasing the drug dose enhanced, rather than suppressed, preference for the drug reward. These studies suggest that the mechanisms controlling choice under drug influence are more complex than previously suggested, in agreement with the present study.

Importantly, motivation for cocaine fluctuates within choice sessions. Indeed, at each cocaine injection, the priming effects of cocaine typically follow a period of lower satiated motivation (Freese et al., 2018; Norman and Tsibulsky, 2006). This satiated motivation is thought to result from a satiating level of dopamine in the ventral striatum (Wise et al. 1995; Ahmed et al. 2003). Indeed, it was shown that a non-contingent injection of cocaine or heroin, which elevates dopamine in the ventral striatum, is sufficient to suppress intra-cerebral self-stimulation of dopamine neurons of the ventral tegmental area (Corre et al., 2018; Pascoli et al., 2015). How drug-induced changes in motivation for the drug and the nondrug rewards interact to dynamically influence preference during choice remains a challenging question deserving further research.

To conclude, our study shows that the mechanisms underlying choice between drug and nondrug rewards under the influence of the drug are complex. A comparable complexity is likely at play when the drug exerts enhancing rather than suppressing effects on the alternative nondrug reward. The present study also reveals that a behavior learned during sequential presentation of the drug and the nondrug rewards (tolerance training) may not generalize well to a choice setting where the drug and nondrug alternatives directly compete with each other for the allocation of behavior. Delineating the complex interactions between motivations for the drug and the alternative nondrug rewards with or without drug influence, is of particular interest for cognitive and behavioral therapeutic approaches.

This research is notably relevant for motivational interviewing aimed at resolving addicted persons’ ambivalence about interacting and competing motivations for their drug of choice and the alternative nondrug-related activities.

## Author contributions

Conceptualization: SHA, YV; Methodology: SHA, YV; Investigation; YV; Formal analysis: YV; Supervision: SHA; Visualization: YV; Writing - original draft: YV; Writing - review & editing: SHA, YV.

