## Supplemental Figures for "Choosing between cocaine and sucrose under the influence: testing the effect of cocaine tolerance"

Vandaele Y.[1], Ahmed S.H[2,3].

[1] Center for psychiatric Neurosciences, Department of Psychiatry, Lausanne University Hospital, Prilly, Switzerland.
[2] Institut des Maladies Neurodégénératives, Université de Bordeaux, Bordeaux, France.
[3] Institut des Maladies Neurodégénératives, CNRS, Bordeaux, France.

Corresponding author: Youna Vandaele
.

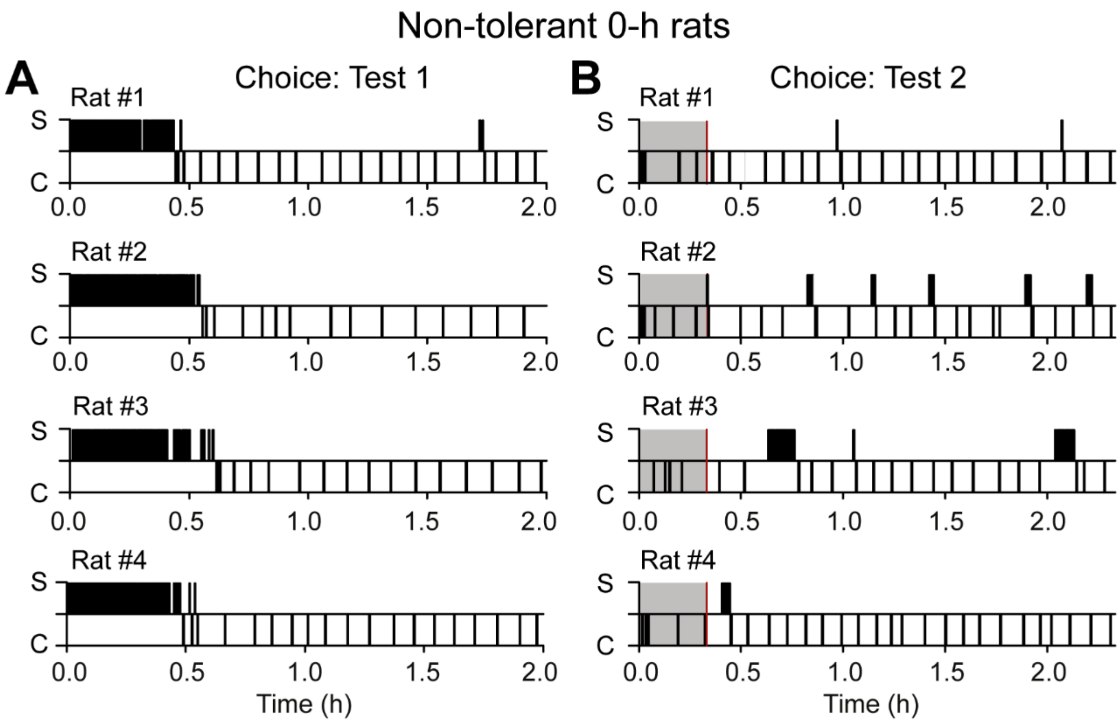


**Supplemental figure 1:** Choice patterns of the four 0-h rats that did not develop a tolerance to cocaine-suppressive effects, in the first (**A**) and second (**B**) choice sessions. Vertical bars above or below the horizontal line represent sucrose (S) and cocaine (C) choices, respectively. In B, the gray area represents the 20-min period of pre-choice cocaine self-administration. The onset of the choice session is marked with a red vertical bar.

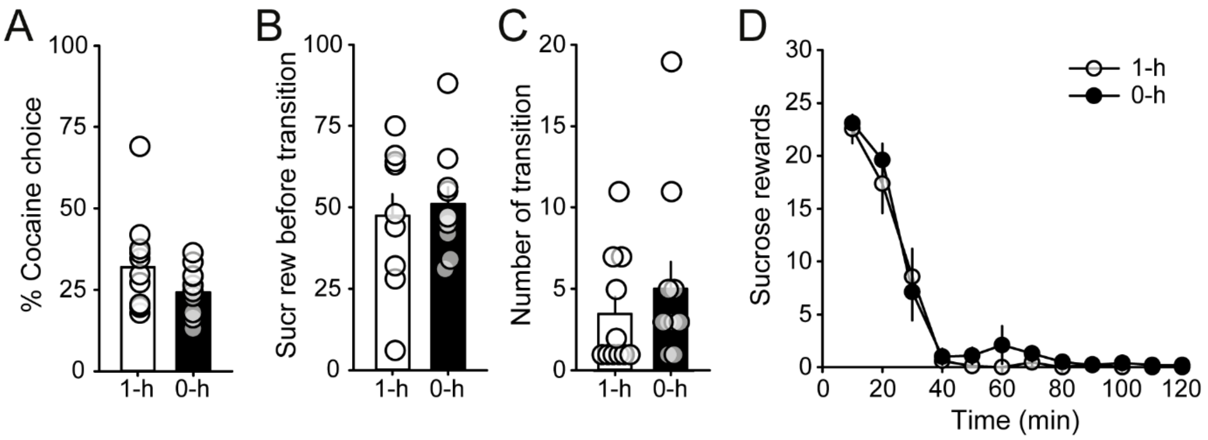


**Supplemental figure 2: Tolerance to cocaine-suppressing effects had no effect on preference during a second 2-h choice session**. **A-C.** Mean (±SEM) percentage of cocaine choice (**A)**, number of sucrose rewards before transition to cocaine (**B**) and number of inter-reward transitions (**C)** in 1-h and 0-h rats during a replication of the 2-h choice session. **D.** Within-session time course of sucrose rewards in the 1-h and 0-h rats across 10-min time bins.
